# Fetal sex is associated with the maternal gut microbiota composition and its metabolic function

**DOI:** 10.1101/2025.06.15.659758

**Authors:** Blake M. Williams, Jessica Liu, Beatriz Peñalver Bernabé

## Abstract

**Background:** Pregnancy affects almost every organ and system, including the microorganisms that resides in the gastrointestinal track, the gut microbiota. Pregnancy hormones, such as progesterone and estradiol, alter the composition of the maternal gut microbiota, directly and indirectly, and in turn, the gastrointestinal microorganism can modify the activity of reproductive hormones. While the levels of pregnancy hormones depend on the fetal sex, the role of fetal sex in the maternal gut microbiota composition and function has not been explored yet.

**Methods:** Using publicly available microbial sequencing data from a cohort of 485 pregnant women in Guangzhou, China, we identified taxonomical and predicted metabolic signatures associated with fetal sex and gestational age using zero-inflated Gaussian models and linear models, respectively, correcting for pre-pregnancy BMI, maternal age, parity, and residency status. Further, we determined sex-specific co-abundance microbial networks using SPIEC-EASI.

**Results:** We observed higher diversity levels in male than in female pregnancies and identified multiple microbial species and predicted microbial enzymes that were differentially abundant between the fetal sex groups, such as increased levels of *Tyzerrella* and β-glucuronidase in female pregnancies, respectively. β-glucuronidase deconjugates glucuronide-bound estrogens into free estrogens and glucuronic acid. Further, the microbial co-abundance networks between the male and female pregnancies were distinct, with efficient information transfer being representative of the female network while the male network was characterized by a larger number of competitive or exclusion interactions between microbial species. While the two networks had similar core communities, there were distinct connections between hormone metabolizing microbes, with β-glucuronidase expressing microbes being more highly connected in the female network.

**Conclusions:** Our study suggests that differences in the maternal-fetal interactions, such as fetus/placenta-produced hormone levels, may affect and be affected by the composition of the gut microbiota of pregnant individuals in a sex-specific manner. A better understanding of the relationship between fetal sex and the maternal gut microbiota may improve maternal health and fetal outcomes.

**Plain English Summary:** The adequate functioning of the microbial communities that reside in the maternal gastrointestinal tract is essential for normal fetal development and their abnormal behavior has been associated with multiple conditions such as preterm birth, gestational diabetes, or depression. Yet, male and female fetuses interact differently with the maternal systems, in part due to the differences in fetal/placental-produced sex hormones. As studies outside pregnancy have shown that estrogen and progesterone metabolites can affect and be affected by the gut microbiota, fetal sex may have direct and indirect effects on maternal outcomes. Understanding these differences could enable better tailoring of prenatal care.

**Highlights:**

- Fetal/placental-produced sex hormones may affect and be affected by the maternal gut microbiota composition and its metabolic functions.
- Gut microbiota diversity and composition during pregnancy is linked to the fetal sex.
- Bacterially produced beta-glucuronidase (GUS), an enzyme that can deconjugate glucuronide-bound estrogens into free glucuronic acid and free estrogen, is increased in pregnancies with female fetuses, plausibly due higher levels of glucuronide-bound estrogens in female pregnancies that benefit GUS-expressing bacterial species that consume glucuronic acid
- The maternal co-abundance gut microbial communities (networks) are distinct in pregnancies with female versus male fetuses specially among taxa that could produce GUS.

## INTRODUCTION

Biological sex plays an essential role in a multitude of biological processes, including cell differentiation, tissue formation, organ functioning, and aging, to name a few^1^. Further, sex differences are common in the prevalence and progression of multiple conditions, including cardiovascular^2^, autoimmune^3^, and neurological disorders^4^, such as multiple sclerosis^5^ and Alzheimer’s disease^6^, and even in the composition and function of the gut microbiota^7^, the microorganisms that reside in the gastrointestinal track^8^. The gut microbiota plays an integral role in human health, including fermentation of non-digestible fibers^9^, production of vitamins^10^, modulation of the intestinal barrier^11^, and immune regulation^12^. Inadequate functioning of the gut microbiota is associated with multiple diseases, including but not limited to inflammatory bowel disease (IBD)^13^, obesity^14^, type 2 diabetes mellitus^15^, autoimmune disorders^16^, neurological conditions^17^, and mood disorders^18^.

The composition and function of the gut microbiota depends on a myriad of factors, including diet, age, environment, geographic location, medication use, social interactions, and biological sex^7,19–24^. Several studies have shown sex-based differences in the gut microbiota structure and composition^7,21,23,25,26^, which could be in part due to different levels of sex hormones. In fact, natural changes in the hormonal milieu, such as menstruation, menopause, and pregnancy lead to alternations in the composition and structure of the gut microbiota^20,27,28^. For instance, increasing levels of progesterone leads to enrichment of *Bifidobacterium* in the third trimester of pregnancy^29^.

Pregnancy is a dynamic period that affects every organ in the body including the immune, endocrine, and nervous systems and their commensal microorganisms. Pregnancy hormones, such as estrogen and progesterone, dramatically increase throughout pregnancy and rapidly decrease after delivery^30^. Circulating hormone levels are fetal sex dependent, with higher testosterone concentrations in maternal blood^31^ and higher levels of 17-hydroxypregnenolone in maternal plasma of male pregnancies. ^32^ The placenta is sexually dimorphic^33,34^ and participates in the production and modulation of pregnancy hormones, such as deconjugation of sulfated steroids into unconjugated active forms ^35^, which may lead to sex-specific differences in maternal and fetal hormone levels. Pregnancy hormones can alter the composition, structure, and function of the gut microbiota both directly and indirectly. Estrogen and progesterone alter the maternal immune system, which is a key regulator of the gut microbiota^36^. Certain microbial species can directly metabolize cholesterol, the precursor for sex steroid hormones, such as *Eubacterium* ATCC 21408 which converts cholesterol to coprostanol, while other species can metabolize sex steroid hormones themselves, including *Eggerthella lenta* 144 which converts androstenedione to testosterone^37^. Additionally, progesterone supplementation both *in vivo* in mice and *in vitro* has been shown to increase the abundance of *Bifidobacterium*^29^. In turn, the gut microbiota can modify the maternal hormonal milieu. For example, microbially produced beta-glucuronidase (GUS) can deconjugate glucuronide-bound estrogens into free estrogens, which can return to circulation through the enterohepatic system, and glucuronic acid^20,38,39^, that bacteria use as energy source. Additionally, gastrointestinal microbes can convert glucocorticoids into progestins, such as allopregnanolone a derivative of progesterone, through 21-dehydroxylation^40^. Further, allopregnanolone, and other bacterially produced progestins are higher in stool samples during pregnancy compared with non-pregnant individuals^40^. Therefore, it is plausible to hypothesize that the sex of the fetus may influence the composition and structure of the maternal gut microbiota and in turn, the gut microbiota has the potential to alter circulating levels of pregnancy hormones. Yet, no study has explored the role of fetal sex in the gut microbiota composition and its metabolic functions.

Here, using a large publicly available dataset of gut microbiota samples during pregnancy, we employed a combination of statistical, computational, and network theory to determine the associations between fetal sex and gut microbiota composition, structure, and metabolic functions. Our results showed that there is indeed a fetal sex-gut microbiota axis during the pregnancy.

## METHODS

### Study cohort and characteristics

We employed a publicly available dataset that include samples from pregnant individuals who were recruited from the obstetric clinics at Guangzhou Women and Children’s Medical Center in Guangzhou, China from January 2017 to September 2017. Sample collection methods and exclusion criteria for the original study cohort are described by Yang et. al^41^. We further excluded participants with complicated pregnancies: i) carrying more than one fetus; ii) taking any kind of medication including progesterone; iii) history of prior and/or current disorders, except for history of allergies or anemia and hepatitis virus infection or immunization. Additionally, we also excluded participants with missing data for fetal sex, pre-pregnancy body-mass index (BMI), residency status (native Chinese or individuals that had resided in Guangzhou for more than a year), gestational age (EGA), parity, and maternal age, rendering a total of 485 participants for downstream analysis.

### Statistical analysis of metadata

F-tests were used to determine statistically significant differences in the variances of continuous phenotypical information based on fetal sex and student’s t-tests and Chi-square tests, respectively, for continuous and categorical information. We employed the following covariates in our models: parity, maternal age, pre-pregnancy BMI, and residency status. Continuous covariates, parity, maternal age, and pre-pregnancy BMI were mean centered and standard deviation scaled.

### Microbiota data analysis and identification of differentially abundant taxa

Publicly available raw 16S rRNA amplicon sequencing data of prenatal stool samples was downloaded from EBI accession number PRJEB31743^41^. Amplicon sequence variants (ASVs) were determined using *DADA2 v1.28.0* in R with standard parameters, unless indicated otherwise^42^. Raw forward and reverse sequences were truncated to 250-256 base pairs, the expected range of read lengths for V4 amplicon region, and denoised, followed by removal of chimeras. ASVs with relative abundance less that 1% were removed from downstream analysis. Taxonomical assignment of denoised and filtered reads was performed using the SILVA reference database version 138.1^43^. Taxonomical assignment and count data were merged using *phyloseq v1.46.0* in R ^44^. ASV count data was normalized using cumulative sum scaling (CSS) ^45^ and then rounded to the nearest integer for downstream analyses.

Alpha diversity was calculated using the Chao1^46^, Shannon ^47^, and Simpson^48^ indices and beta diversity was estimated using Bray-Curtis dissimilarity distance^49,50^. Multivariable linear regression was used to model the changes in alpha diversity indices by fetal sex, EGA, and their interaction. Fetal sex and EGA were included as factors in all models throughout the analysis and models were corrected for pre-pregnancy BMI, maternal age, residency status and parity. Residency status was included in the model as it can be a proxy for different type of diets or cultural habits, which are known to affect the gut microbiota composition and metabolic function^24^. To remove the effect of covariates, we calculated the covariates-corrected alpha diversity distance, *X̂_ij_* as follows:

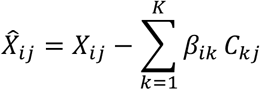

where *X_ij_* is the alpha diversity distance, *β_ik_* is the estimated effect (beta coefficient) of covariate *k* on alpha diversity distance *i*, and *C_kj_* is the value of covariate *k* for sample *j*. We then used unpaired Wilcoxon rank-sum test^51^ with exact p-value estimation to determine statistically significant differences in the fitted alpha diversity metrics (Chao1, Shannon, and Simpson) by fetal sex. Statistically significant differences in beta diversity between the fetal sex groups were assessed using PERMANOVA^52^, correcting for the same covariates as above.

Zero-inflated Gaussian models^29^ were employed to identify differentially abundant ASVs by fetal sex, EGA, and their interaction, correcting for pre-pregnancy BMI, maternal age, residency status and parity with the *metagenomeSeq v1.42.0* package in R^45^. All models were corrected for multiple comparisons using the false discovery rate (FDR) method (FDR adjusted p-value < 10^−3^). ASVs were agglomerated at the lowest identified taxonomic level prior to plotting.

### Predicted microbial metabolic potential

Bacterial metabolic potential was predicted from CSS-normalized ASV count data using *PICRUSt2 v.2.5.2* ^53–57^. CSS-normalized ASVs and their associated denoised FASTQ sequences were input into the *PICRUSt2* pipeline to predict bacterial enzyme counts. Predicted bacterial enzymes with minimum prevalence of 90% and abundance in less than 90% of samples were removed from downstream analysis, and subsequently they were CLR transformed. Using *MaAsLin2 v1.14.1* ^58^, we built multi-linear regression models to identify statistically significant differentially predicted bacterial enzymes in terms of their abundance as a function of fetal sex and EGA, including their interaction term. We included all the covariates listed above and corrected for multiple comparisons (FDR-adjusted p-value < 0.25). We repeated this analysis using *PICRUSt2* predicted MetaCyc pathways^59^ as described above. Functional enrichment analysis was performed in *MicrobiomeProfiler v1.6. 1* ^60,61^ in R using the subset of Kyoto Encyclopedia of Genes and Genomes (KEGG)^62,63^ enzymes that was identified as significantly associated with fetal sex (unadjusted p-value < 0.1).

### Microbial co-abundance network analysis

Microbial co-abundance networks for each fetal sex were inferred from CSS-normalized ASV count data using *SPIEC-EASI v1.1.3* ^64^. ASVs included in the analysis were the top 100 most abundant ASVs and the statistically significant differentially abundant ASVs by fetal sex, EGA, and their interaction (n=21). Statistically significant edges for each fetal sex co-abundance network were identified using bootstrapping. We generated a total of 100 co-abundance networks with *SPIEC-EASI* using the glasso method with *lambda.min.ratio* of 10^−2^ and *nlambda* of 20 using a randomly selected sample of 80% of the participants in each bootstrapping iteration. Null models for each of the 100 iterations were created by randomly shuffling the count data between samples of each ASV to preserve the distribution of each ASV. Edges with an FDR-adjusted p-value < 10^−10^ were deemed significant against the null model using a paired t-test. After removal of non-significant edges, we determined multiple local and global centrality measurements using NetworkAnalyzer^65^ in Cytoscape^66^ assuming edges were unweighted and undirected: a) average shortest path length (measurement of the efficiency of information transfer within a network, the average number of steps along the shortest path from each node to every other node in a network^67^); b) clustering coefficient (measurement of how closely connected nodes are within a network or the likelihood that nodes within a network will cluster together^68^); c) closeness centrality (the reciprocal of the sum of the shortest path lengths for one node to all other nodes in the graph^67,69^); d) eccentricity (the maximum distance between a node and all other nodes^70^); e) stress (number of shortest paths traversing through a given node^69^); f) degree (number of edges of a given node); and g) edge betweenness (number of shortest paths through an edge^69^). Wilcoxon signed-rank test^71^ and Wilcoxon rank sum test^51^ were used to determine differences between global centrality measurements and edge betweenness, respectively, for the male and female networks. Students t-statistic tests were used to determine the statistically significant was calculated for the global and local centrality measurements, with p-value < 0.01 deemed significant.

### Predicting fetal sex from microbial taxa and microbial metabolic potential

Three machine learning models were utilized to predict fetal sex: Random Forest and Logistic Regression from the *scikit_learn v1.5.0* package^72^ and XGBoost from the *XGBoost v2.0.3* package^73^ in Python. We employed four different input data for the models: (1) biometric data including EGA, maternal age, pre-pregnancy BMI, parity, and residency status; (2) 36 ASVs significantly associated with fetal sex (unadjusted p-value < 0.05) and biometric data; (3) 45 predicted microbial metabolic potential significantly associated with fetal sex (unadjusted p-value < 0.05) and biometric data; and (4) above datasets, biometric data, ASV counts, and predicted microbial metabolic potential, combined. Datasets 2 and 3 were CLR normalized^74^ independently, then combined to create dataset 4. Continuous metadata variables were centered and scaled between 0 and 1 for all datasets. Hyperparameter tuning was performed using nested k-fold cross validation, with ten outer folds for testing and five inner folds for hyperparameter tuning, splitting the data 80 to 20 for training and testing sets, respectively, for each fold. Using *GridSearchCV* from *Scikit-learn v1.5.0* in Python^72^, we tuned each model’s parameters.

## Results

### Gut maternal microbial composition differed by fetal sex

We used publicly available 16S rRNA amplicon sequencing data of maternal fecal samples from a cohort of xxx pregnant individuals from a clinic in Guangzhou, China. After removing participants with clinical data that might affect the composition or structure of the gut microbiota (e.g., pregnancy complications, medication), we selected a total of 485 participants (female fetus: 239, male fetus: 246, **Figure 1, Table S1)**. Most participants were native Chinese born in the city of Guangzhou (female fetus: 70.3%; male fetus: 72.0%). The rest of participants were Chinese individuals not native to Guangzhou but who had resided there for at least one year prior to enrolling in the study. Most of the donated samples were in the second trimester of gestation (57%, 14-27 weeks), while 7.4% and 35.1% were collected during the first (<13 weeks) and third (28-40 weeks) trimesters, respectively. We didn’t observe any statistical differences by fetal sex in terms of maternal age, estimated gestational age (EGA), pre-pregnancy BMI, parity, or residency status (p > 0.05, **Table 1**).

**Figure 1:**
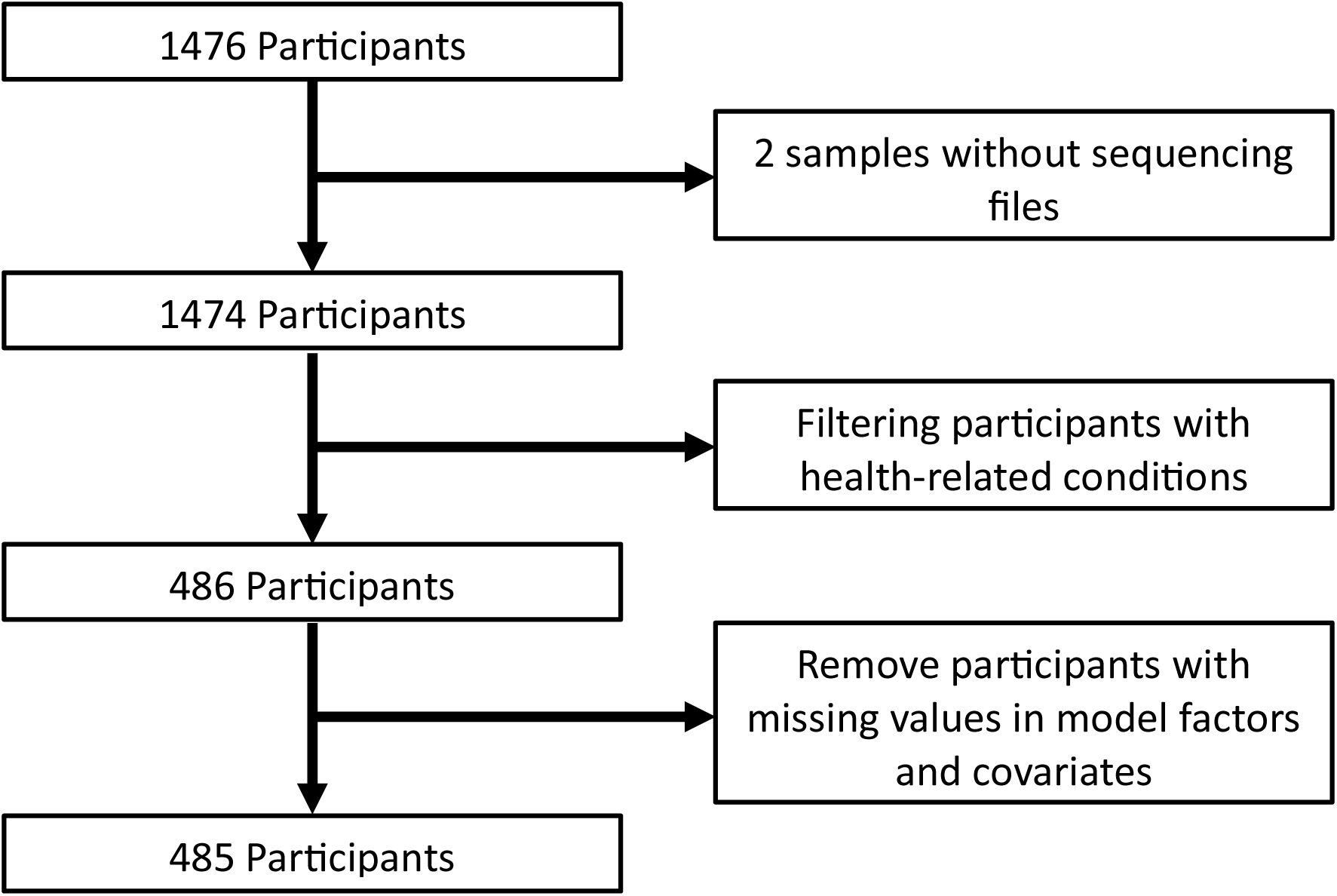
Participant filtering schematic. Data from original study was filtered sequentially as shown in schematic.

**Figure 2:**
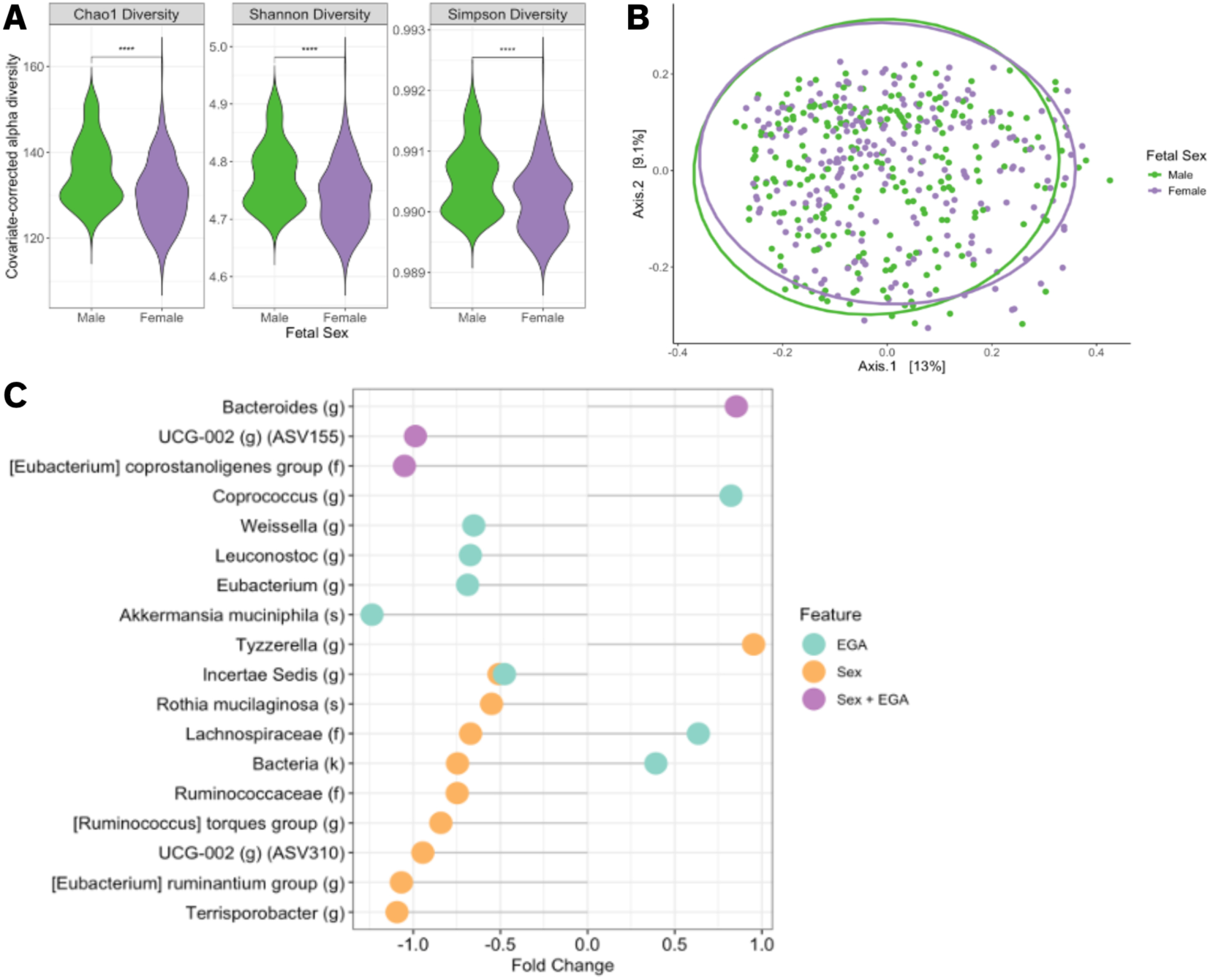
Microbial diversity was associated with fetal sex. **(A)** Alpha diversity showed significant differences for male and female fetuses. Alpha diversity was calculated using Chao1, Shannon, or Simpson indices. Multilinear regression was used to model alpha diversity by sex correcting for maternal age, residency status, pre-pregnancy BMI, parity, and the interaction between fetal sex and EGA. Sample size: 485 pregnant individuals. Significant differences were also observed for fetal sex (****p<1×10^−15^) via Wilcoxon rank-sum test. **(B)** Beta diversity, calculated using Bray-Curtis dissimilarity, showed no statistically significant differences in community structure by fetal sex, but significant differences in by residency status (p < 0.001) and parity (p < 0.05) via PERMANOVA analysis. **(C)** Several microbial species were associated with fetal sex, EGA, and their interaction. Significant differentially abundant microbial species were assessed using zero-inflated Gaussian Models, correcting for residency status, parity, maternal age, and pre-pregnancy BMI. Microbial taxa increased or depleted in individuals pregnant with female fetuses (orange), EGA (teal), and their interaction (purple) with FDR-adjusted p-value < 10^-3^.

**Table 1:** Sample metadata and model covariates. There were no statistically significant differences between pregnancies carrying female and male fetuses for the metadata covariates included in our models. Continuous variables are reported as mean (sd); binary variables are reported as count (percent). Student’s t-test for continuous variables and Chi-square for categorical variables.

|  | N (Percentage) |  |
| --- | --- | --- |
|  | Female (N=239, 49%) | Male (N=246, 51%) |
| Residency status | 168 (70.3%) | 177 (72.0%) |
|  | Mean (sd.) |  |
| Maternal age | 29.9 (3.92) | 29.7 (3.83) |
| Gestational weeks | 23.3 (7.45) | 23.9 (7.38) |
| Pre-pregnancy BMI | 20.4 (2.37) | 20.1 (2.39) |
| Parity | 0.39 (0.51) | 0.41 (0.52) |

We then contrasted the gut microbiota structure and composition as a function of fetal sex. While there were no statistically significant differences in alpha diversity by fetal sex for the uncorrected Chao1, Shannon, and Simpson indices (**Figure S1,** p > 0.01), we did uncover statistically significant differences by fetal sex after correcting for maternal age, EGA, pre-pregnancy BMI, residency status, and parity using multilinear regression models (p < 0.01, **Fig. 1A**). For beta diversity, as measured by Bray-Curtis dissimilarity distance, we did not observe any statistically significant differences by fetal sex using PERMANOVA (**Fig. 1B)**, but did find significant differences by residency status (p < 0.001) and parity (p < 0.05).

At the taxonomical level, our results indicated that the composition of the gut microbiota was dependent on fetal sex, EGA, and their interaction (**Fig. 1C**). We identified a total of ten ASVs that were significantly associated with fetal sex (e.g., unspecified species of the genera *Terrisporobacter* and *Tyzzerella*), eight associated with EGA (e.g., *Akkermansia muciniphila* and unspecified species of the genus *Leuconostoc*), and three associated with their interaction (e.g., unspecified species of the genera *Eubacterium coprostanoligenes* group and *UCG-002*).

Independently of the gestational age, only an unclassified species from the *Tyzzerella* genus was increased in female pregnancies, while *Rothia mucilaginosa*, nine species from the Clostridia class, and one uncharacterized species were positively associated with male fetuses. An uncharacterized species from the *Incertae Sedis* genus was independently associated with both sex (higher in male pregnancies) and EGA (lower as pregnancy progresses). Importantly, we identified several taxa whose association with fetal sex depends on the gestational period. For instance, an unclassified member of the *Bacteroides* genus increased in female pregnancies as gestation progressed while an uncharacterized species from the *Eubacterium coprostanoligenes group* decreased in female pregnancies as gestation progress. Overall, we found significant differences in gut microbial diversity and composition between the fetal sex groups, yet some of those associations were dependent of the gestational period and thus, likely, on the total level of circulating pregnancy hormones. ^28,38^

### Predicted metabolic potential of the gut microbiota was associated with pregnancy hormone metabolism

Using *PICRUSt2* ^53–57^ to predict the most likely genes that encode for enzymes given the ASVs identified in the samples, we identified several statistically significant differences in the bacterial metabolic potential as a function of fetal sex (**Figure 3A)**. Specifically, eight predicted bacterial enzymes were higher in pregnancies with female fetus, such as starch synthase (glgE), bacterial fatty acid synthase (fas), dextranase (dexA), and putative RNA 2’-phosphotransferase (kptA) and four were lower, including flavin phenyltransferase (ubiX), gamma-glutamyltranspeptidase/glutathione hydrolase (ggt), 1,4-dihydroxy-2-napthoyl-CoA hydrolase (menI), and diguanylate cyclase (adrA). Of the eight enzymes higher in female pregnancies, shikimate kinase/3-dehydroquinate synthase (aroKB) also increased as gestation progressed. Additionally, three proteins related to the glutamate transport system, including the substrate binding protein (glub) and two permease proteins (gluC and gluD), increased in female pregnancies and as gestation progressed. Of note, beta-glucuronidase (uidA or microbial GUS), a bacterial enzyme that can deconjugate glucuronide-bound estrogens to produce free, biologically active estrogens^38,39^, was significantly enriched in pregnancies with female fetuses independently of their gestational age. We also identified multiple enzymes that were enriched in male pregnancies, such as gamma-glutamyltranspeptidase/glutathione hydrolase (ggt) and purine/pyrimidine-nucleoside phosphorylase (ppnP) and the transcriptional regulator, CopG family transcriptional regulator/antitoxin EndoAI (ndoAI).. Finally, a multitude of predicted microbial enzymes(?)were associated with EGA (**Table S3**) andfour of them their associations with gestational period were dependent on the fetal sex, including galacturonosyltransferase WbtD (wbtD), two-component system NtrC family sensor histidine kinase AtoS (atoS), sugar phosphatase (ybiV), and mannose-1-phosphate guanylyltransferase/phosphomannomutase (K16881). Further, we found that many enzymes associated with fetal sex had different response patterns as pregnancy progressed, such as beta-glucuronidase which was consistently higher female pregnancies but decreased over time in male pregnancies with the largest decrease toward the end of gestation (**Figure 3B**). In terms of metabolic pathways (MetaCyc pathways)^59^ we didn’t find any to be significantly associated with fetal sex.

**Figure 3:**
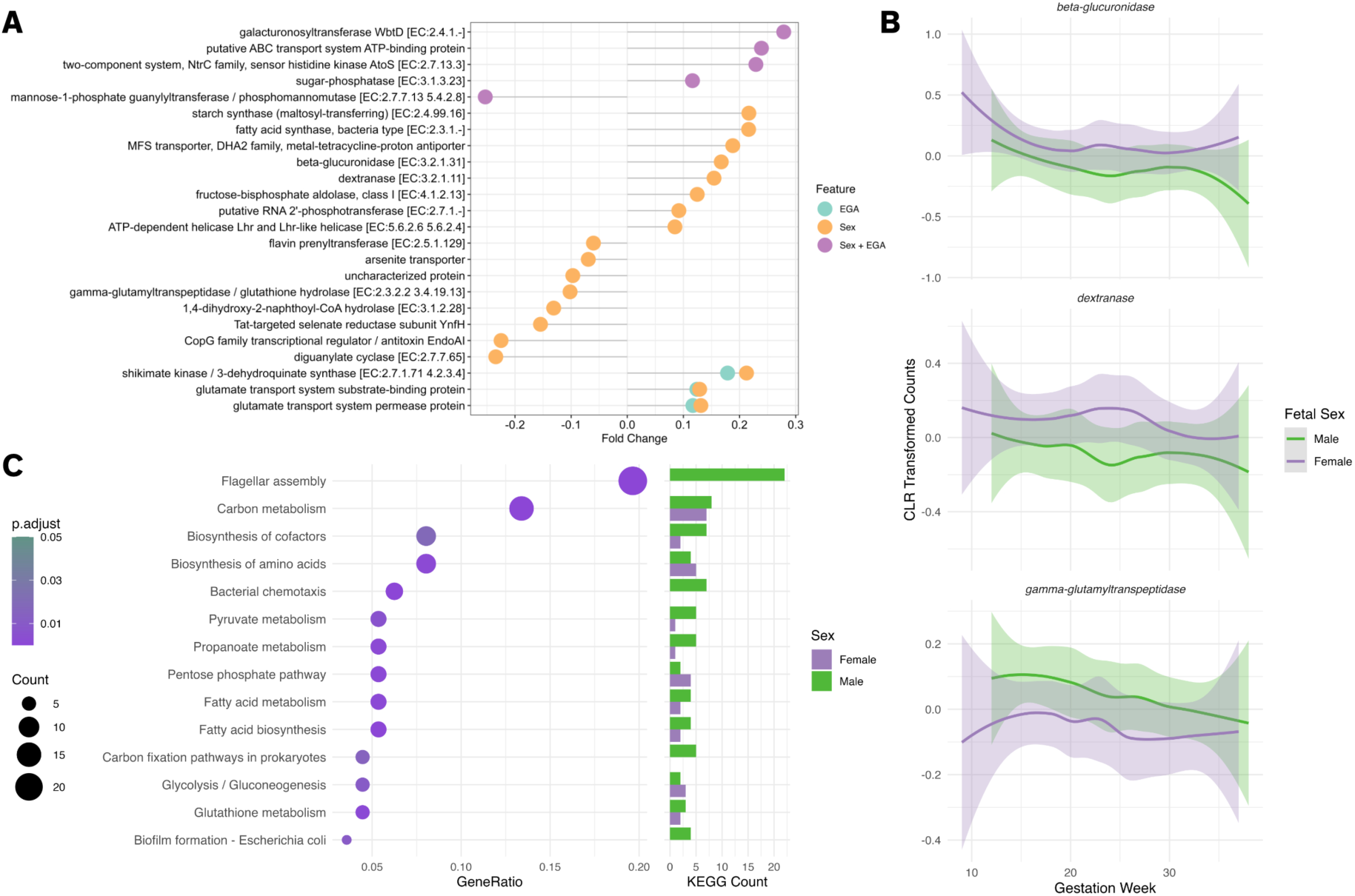
Bacterial metabolic potential was associated with fetal sex. **(A)** Several predicted bacterial enzymes were associated with fetal sex, EGA, and their interaction. Significant enzymatic differences were assessed using linear models correcting for residency status, parity, maternal age, and pre-pregnancy BMI. Enzymes increased or depleted in individuals pregnant with female fetuses (orange), EGA (teal), and their interaction (purple) with FDR-adjusted p-value < 0.25. **(B)** Temporal trends of selected fetal sex-associated enzymes across gestation with a smooth curve based on CLR-transformed count data. **(C)** Functional analysis using Gene Set Enrichment analysis of significant sex-associated KEGGs (p-value < 0.1) with bar plot display the number of enzymes increase in female (purple) and male (green) for each pathway with FDR-adjusted p-value < 0.05.

We then asked whether those metabolic significant features were enriched for certain metabolic pathways. In total, we identified fourteen KEGG metabolic pathways, (**Figure 3C**; FDR-adjusted p-value < 0.05) many of which were associated with energy production, including carbon metabolism, pyruvate metabolism, and glycolysis/gluconeogenesis, and multiple KEGG metabolic pathways were equally enriched in both male and female pregnancies, such as glutathione metabolism, fatty acid biosynthesis and metabolism, and biosynthesis of cofactors and of amino acids. A total of four pathways were enriched in enzymes whose predicter abundances were higher in pregnancies with male fetuses, including flagellar assembly, bacterial chemotaxis, biofilm formation – *E. coli*, and carbon fixation pathways in prokaryotes (**Figure 3C**). In contrast, while no pathways only contained enzymes enriched in female pregnancies, the energy-relevant pentose phosphate pathway was enriched in mostly female-associated enzymes (n=4/6).

Finally, we explored whether our models were successful at predicting fetal sex for our cohort using multiple machine learning methods, including Random Forest, Logistic Regression, and XGBoost. All the models have a very low predicted capability (AUC<0.6) to determine fetal sex using microbial taxa, predicted bacterial metabolic potential, or a combination of the two (**Fig. S4**). In summary, our results suggested that there are key differences in bacterial composition and metabolic potential in the maternal gut of individuals pregnant with male and female fetuses, but those differences are not predictive of the fetal sex.

### Co-abundance microbial networks differed by fetal sex

As microorganisms do not live in isolation, we asked next whether co-abundance microbial networks were distinct by fetal sex (**Figure 4)**. Using *SPIEC-EASI*^64^, we observed that in the male co-abundance microbial network there were more edges overall connecting different ASVs and greater proportion of negative connections between those ASVs than those of female pregnancies (**Table 2**, p < 0.01), suggesting a greater levels of exclusion and competitive environment in the microbial communities of individuals carrying male fetuses. Male co-abundance networks were characterized by higher closeness centrality (a measurement of how easily a node can reach other nodes) and higher edge betweenness (a measurement of the importance of an edge based on the number of shortest paths that traverse through it) compared with the female networks, while female co-abundance microbial network contained several disconnected small subnetworks (**Table S5, S6**).

**Figure 4:**
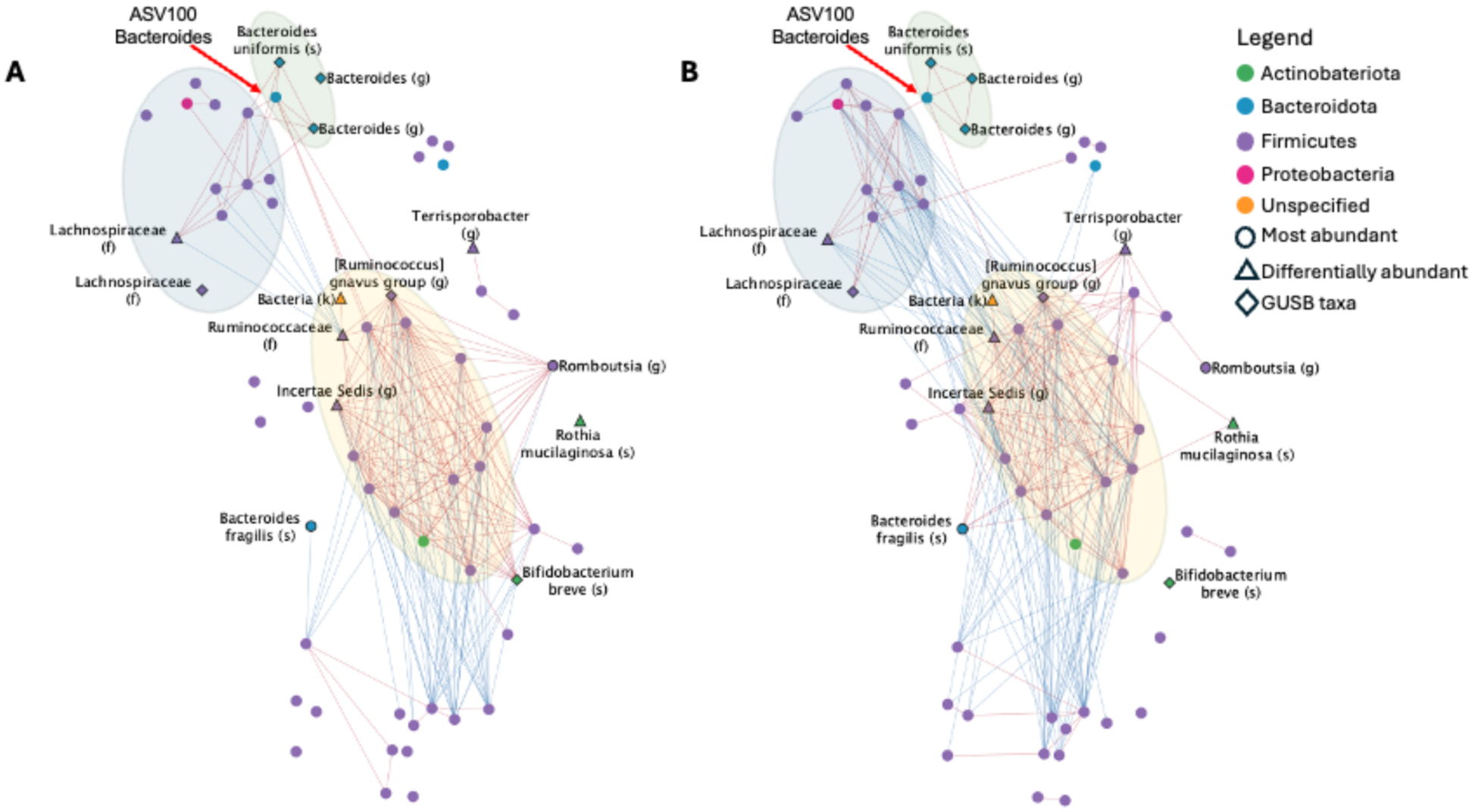
Co-abundance microbial networks were distinct in pregnancies carrying female (A) and male (B) fetuses. Co-abundance networks were obtained using SpiecEasi. Significant edges were identified using bootstrapping (p<10E-10, see methods for more details). Edge thickness is proportional to the edge covariance and color corresponds to sign of the interaction (positive: red; negative: blue). Significant taxa (triangles) were assessed using linear models. Most abundant taxa (circles) were identified from CSS normalized counts. Taxa predicted to encode beta-glucuronidase (diamonds) via PICRUSt2.

**Table 2:** Co-abundance microbial network properties. Comparisons between of node and edge properties and percentage of negative edges for microbial network of individuals pregnant with female and male fetuses were performed via Wilcoxon signed-rank test (node), Wilcoxon rank sum test (edge), and Pearson’s Chi-squared test (percent negative edges), respectively. Values are input as mean (standard deviation). Ns, not sifnificant

|  |  | Mean (sd.) |  | p-value |
| --- | --- | --- | --- | --- |
|  |  | Female (N=239) | Male (N=243) |  |
| Node Properties | Average Shortest Path Length | 1.7 (1.2) | 1.9 (0.8) | ns |
|  | Clustering Coefficient | 0.42 (0.44) | 0.55 (0.39) | ns |
|  | Closeness Centrality | 0.37 (0.28) | 0.53 (0.24) | 5.5E-04 |
|  | Eccentricity | 3.6 (2.6) | 3.1 (1.3) | ns |
|  | Stress | 170 (360) | 290 (450) | ns |
|  | Degree | 6.6 (7.6) | 9.0 (8.7) | ns |
|  | Betweenness Centrality | 0.04 (0.1) | 0.02 (0.03) | ns |
| Edge Properties | Edge Betweenness | 20.1 (34.1) | 20.4 (25.0) | 2.5E-08 |
| Percent Negative Edges |  |  |  | 6.5E-03 |

Both male and female co-abundance networks presented a large well-connected central community (highlighted in yellow) containing a large proportion of positive edges between many microbes, indicating consistent cooperation between microbes in this community regardless of fetal sex. Yet, there were several differences in the network structures. For instance, while there was some weak competition between a few microbes in the female network (the blue and yellow communities), the competition between these two communities was much stronger in the male network as demonstrated by the greater number of negative interactions between ASVs (blue edges). Additionally, the green community consisting of primary of GUS-expressing (diamond-shaped nodes) *Bacteroides* species had many cooperative connections to two other communities in the female network (blue and yellow subcommunities). Yet in the male network, one non-GUS-expressing *Bacteroides* node, ASV100 (red arrow), is mediating the interaction between the blue, yellow, and green communities.

Finally, we explore whether the role of certain taxa within the co-abundance networks change as a function of fetal sex (**Table S9**). We observed that all hormone-metabolizing and differentially abundant species had multiple statistically significant centrality measurements as a function of fetal sex. For instance, *Bifidobacterium breve,* a microorganism that can produce GUS (diamond-shaped nodes), had significantly lower average shortest path length, closeness centrality, clustering coefficient, eccentricity, and stress centrality in the female network than the male network (all p-values < 0.01), indicating that *B. breve* was a central hub in the female network. In fact, *B. breve* in not connected to the core community in the male pregnancies. On the other hand, *Bacteroides fragilis* a species that metabolizes 11β-hydroxyandrostenedione and androstenedione into 11β-hydroxytestosterone and testosterone, respectively, through 17β-hydroxysteroid dehydrogenase (17β-HSDH)^37^, had significantly higher betweenness centrality, clustering coefficient, and stress centrality and lower average shortest path length and eccentricity in the male than in the female co-abundance network (all p-values < 0.01). *B. fragilis* was more connected in the male network, with multiple interactions with the core (yellow) community while *B. fragilis* became peripherical in the female co-abundance network. Several ASVs that were significantly associated with fetal sex (triangle-shaped nodes) were also key elements in the co-abundance microbial networks. For instance, many species that were increased in male pregnancies were also more connected in the male co-abundance network, including ASVs mapped to *Rothia mucilaginosa*, unclassified species of the genus *Terrisporobacter*, *Lachnospiraceae*, *Ruminococcaceae*, and *Incertae sedis*. Of these, both *R. mucilaginosa* and the unclassified species of *Terrisporobacter* had positive associations with the core community (yellow) and played a more central roles in the male network as measured by closeness centrality, clustering coefficient, and eccentricity centrality (p-value < 0.01) than in the female co-abundance networks. Interestingly, an ASV assigned to the genera *Ruminococacceae* had the highest stress in both networks, suggesting that it is integral for maintaining network connectivity irrespective of fetal sex. Taken together, these differences suggest that microbial communities were indeed distinct for pregnancies with male and female fetuses, with male pregnancies characterized by more overall interconnected communities with high levels of possible competition and exclusion mechanisms compared with female pregnancies. Further, estrogen- and testosterone-metabolizing microbes played central roles in the maternal gut microbial communities.

## Discussion

Our study demonstrates, for the first time, that the gut microbiota composition, structure, and potential metabolic function are significantly associated with fetal sex. Using publicly available data from a pregnant cohort, we found significant differences in the composition of the maternal gut microbiota for individuals pregnant with female versus male fetuses and we identified multiple associations between fetal sex and taxonomical, metabolic, and community features. These results highlight the importance of including fetal sex as a biological variable when studying the perinatal microbiome.

While similarities in the core community of the co-abundance microbial networks were expected between pregnancies carrying male and female fetus, as conserved pregnancy-related changes occur irrespective of fetal sex, differences in these co-abundance microbial networks underscores the fetal sex’s direct and indirect role in regulating maternal microbial communication. The co-abundance microbial networks in male pregnancies had more overall connections and significantly more negative associations between microbial species. While the current data doesn’t allow us to differentiate, negative connections between microorganisms might be due to competition for similar resources or to the production of toxin components for other members in the community (exclusion), while positive interactions may be due to cooperation or synergistic processes between species. In other words, male pregnancies were characterized by more competitive gut microbial communities, while female pregnancies, although less connected, presented a more cooperative behavior. However, the communication between the microorganisms in the male networks is possibly more efficient, as underscored by their higher closeness centrality and edge betweenness, suggesting a resilient mechanism against competing and decremental species.

Microbial GUS is a bacterial enzyme that deconjugates glucuronide-bound estrogens rendering bioactive free estrogens and free glucuronic acid^20,38,39^. Free glucuronic acid is used as an energy source by the GUS-encoding bacteria^75^ and the free estrogens can enter back into the maternal blood stream through the enterohepatic circulation system and thus bind to estrogen receptors (ER) in human cells^38,39^. We found that the predicted levels of microbially-produced GUS were higher in pregnancies with female fetuses, and that *B. breve*, a GUS-producing species, was a central hub in the female co-abundance network yet it did not play a role in the male community.

Recent studies have ^2931^shown sex differences in hormone signaling and metabolic functions in the placenta, with higher levels of steroid sulfatase (STS), an enzyme that converts sulfated steroid precursors to active free steroids, in female placentas, as well as higher levels of cytochrome P450 subfamily 11 alpha 1 (CYP11A1), which converts cholesterol to pregnanolone (a precursors for all androgens and estrogens) in first trimester male placentas^76^. Additionally, female fetal livers have a higher expression of *UGT2B17*, a gene encoding an enzyme that conjugates free estrogens to glucuronic acid, when compared to male fetal livers^77^. Therefore, it is plausible that sex differences in the placenta and liver functions in the production of glucuronide-bound estrogens might promote the growth of GUS-producing bacteria in female pregnancies as these microorganisms use them as energy source and in turn lead to higher levels of circulating free estrogens (**Figure 5**). Estrogens regulate the immune response by controlling the innate and adaptive immune reactions^78^ and might explain why female fetuses are less susceptible to maternal inflammation^79,80^ and reduced pregnancy complications in female pregnancies in general^81^. In line with this hypothesis, multiple taxonomical species that were more abundant in male pregnancies, such as *Terrisporobacter*, *Rothia mucilaginosa* and *Incertae sedis*, were also highly interconnected in the male networks suggesting potential increased crosstalk between these species that enable them to thrive together in male pregnancies, have been associated with complications in pregnancies carrying a male. For instance, carrying a male fetus increases the relative risk of GDM compared with carrying a female fetus^82^ and individuals with GDM during their second trimester of pregnancy present elevated levels of *Incertae sedis* compared with non-GDM pregnancy^83^. It is tempting to suggest that the lower risk of prenatal complications in female pregnancies might be due, in part, to the prenatal adaption of the gut microbiota.

**Figure 5:**
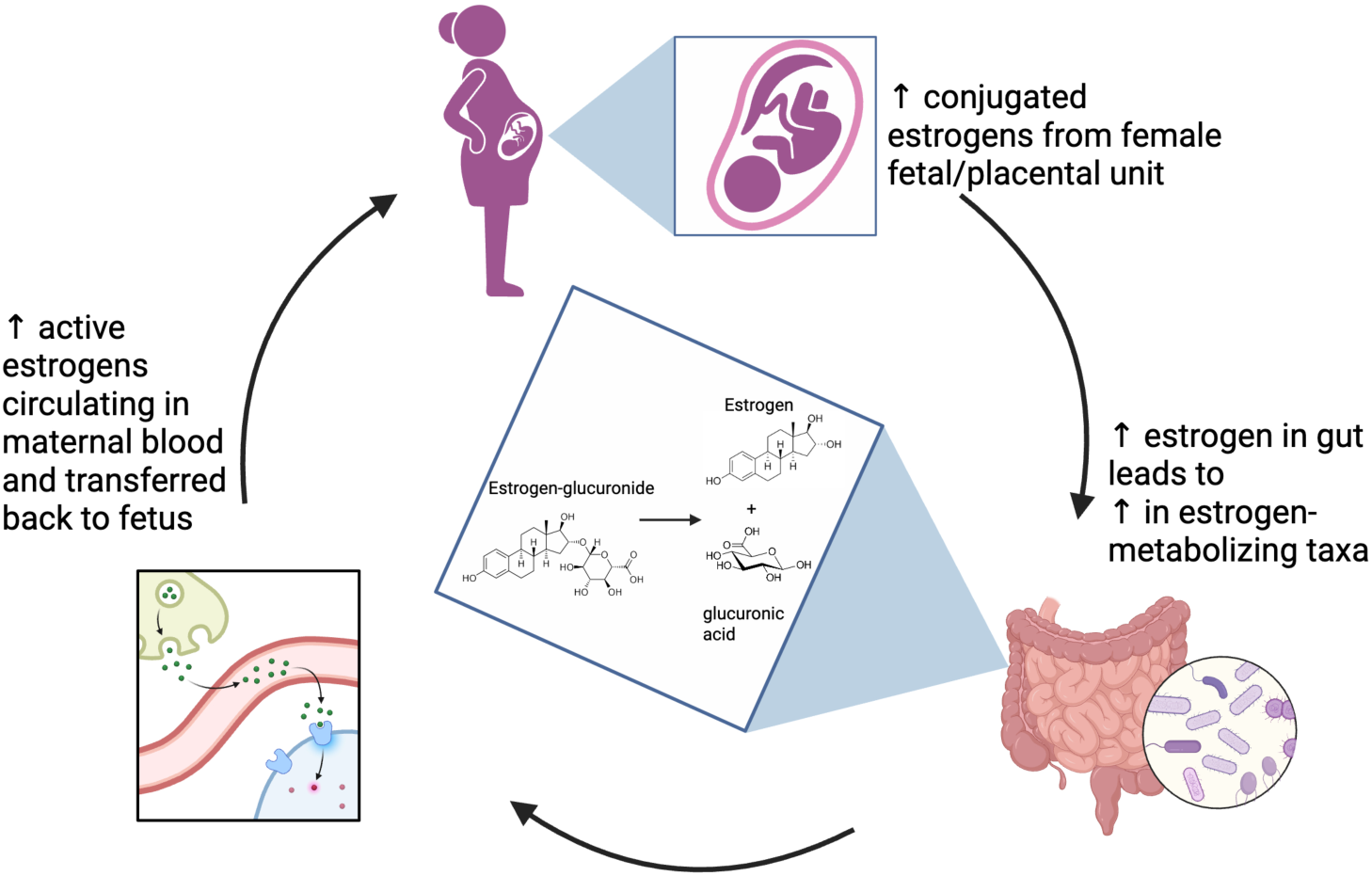
Working hypothesis: Fetal/placental estrogens effect maternal gut microbiota. Increasing levels of fetal/placental-produced estrogens might enrich beta-glucuronidase-expressing taxa (*Bacteroides*, *Bifidobacterium*, and *Ruminococcus*). Bacteria that produce beta-glucuronidase can deconjugate glucuronic-bond estrogens increasing circulating levels of free estrogens which can be transferred back to the mother and the growing fetus through maternal blood.

Meulenberg et al. has shown that maternal testosterone levels are higher during the second half of gestation for male pregnancies^31^ and these higher testosterone levels might be partially explained by the maternal gut microbiota. For instance, *Bacteroides fragilis,* which was more abundant in male pregnancies and was also one of the key members of the co-abundance male microbial communities, metabolizes 11β-hydroxyandrostenedione and androstenedione into 11β-hydroxytestosterone and testosterone, respectively, through 17β-HSDH^37^.

Some microorganisms might also take advantage of the prenatal hormonal enriched environment. As recently reviewed by Ridlon et al., there are multiple paths in the axis between the sex steroid hormones and gut microbiota ^85^. Progesterone leads to proliferation of *Bifidobacterium breve* and while the mechanism is not clear, Nuriel-Ohayon et al. suggested that *Bifidobacterium breve* can produce hydroxysteroid dehydrogenase and thus metabolize progesterone ^29^. Additionally, several microbially-produced enzymes in commensal gut bacteria can metabolize cholesterol, the precursor of sex steroid hormones, as well as sex steroid hormones themselves^37^. Apart from 17β-HSDH which converts androstenedione to testosterone, 16-dehydroxylase metabolizes 16α-hydroxyprogesterone to render 17α-pregnanolone and 17α-hydroxysteroid dehydrogenase (17α-HSDH) transforms androstenedione to epitestosterone^37^. Additionally, *Gordonibacter pamelaea* and *Eggerthella lenta* metabolize biliary corticoids into progestins, such as allopregnanolone^40^, a neuroactive steroid recently approved to treat postpartum depression^85^. It is plausible that these microbially-produced enzymes present different activity in the maternal gut of male and female pregnancies. However, we couldn’t confirm this hypothesis as none of these enzymes were included in the current *PICRUSt2 v.2.5.2* database, and further research is needed.

To the best of our knowledge, this is the first to study to explore the association between fetal sex and the maternal gut microbiota. While the sample size that we employed was large, the study was cross-sectional. As the gut microbiota is unique in each individual^86^ and evolves during pregnancy^28^, it is necessary to perform prospective longitudinal studies to understand the role of maternal microbiota in perinatal outcomes. Further, this study relies on 16S rRNA amplicon sequencing, which does not provide information about gene expression or metabolic potential^87^. Future studies that leverage metagenomics, metatranscriptomics or metaproteomics and metabolomics are necessary to provide a better understanding of the functional characteristics of the maternal microbial communities. Finally, as diet is one of the main regulators of the gut microbiota ^24^, studies that include participants from different regions of the world and with varied food intake profiles are necessary to conclude whether these sex-specific differences in predicted microbial metabolic potential are universal or cohort specific.

## Conclusion

Overall, our study underscores the potential role of fetal sex on the maternal gut microbiota and its metabolic functions. Our results highlight that differences in fetal/placental-produced hormone levels may affect and be affected by the composition of the gut microbiota of pregnant individuals in a sex-specific manner, yetFurther research is necessary to elucidate the bidirectional relationship between pregnancy hormones and maternal microbiota. A better understanding of how fetal sex regulates maternal gastrointestinal microbial communities, which are key for a healthy pregnancy, could aid to mitigate pregnancy-related complications and thus protect the health of the growing fetus and the mother.

## Acknowledgments, contributions

We would like to thank Dr. Shenghui Li and colleges, the researchers that collected the data that our study relays upon ^41^, for their assistance. We would also like to thank Dr. Melissa Wilson for her curiosity that inspired this study. BPB and BW designed the study and wrote the first draft of the manuscript. BW and JL performed the statistical and computational analysis. All the authors analyzed and interpret the results and reviewed the final manuscript.

## Data and code availability

Raw FASTQ files are available in the European Nucleotide Archive (ENA) study accession number PRJEB31743 and updated sample names are detailed in **Table S10.**

## Code availability

Code is available at https://github.com/LabBea/sex_diff_microbiota

